# Plasticity versus stability across the human cortical visual connectome

**DOI:** 10.1101/520395

**Authors:** Koen V. Haak, Christian F. Beckmann

## Abstract

Whether and how the balance between plasticity and stability varies across the brain is an important open question. Within a processing hierarchy, it is thought that plasticity is increased at higher levels of cortical processing, but direct quantitative comparisons between low- and high-level plasticity have not been made so far. Here, we addressed this issue for the human cortical visual system. By quantifying plasticity as the complement of the heritability of functional connectivity, we demonstrate a non-monotonic relationship between plasticity and hierarchical level, such that plasticity decreases from early to mid-level cortex, and then increases further of the visual hierarchy. This non-monotonic relationship argues against recent theory that the balance between plasticity and stability is governed by the costs of the “coding-catastrophe”, and can be explained by a concurrent decline of short-term adaptation and rise of long-term plasticity up the visual processing hierarchy.

## Introduction

Cortical plasticity, the reorganization of neural circuits in response to environmental change, is ubiquitous in the brain across the lifespan. Cortical plasticity is typically considered to be beneficial because it optimizes neural processing in the face of changing environmental conditions, injury and disease. Within processing hierarchies, however, excessive plasticity at one level of processing could disrupt the functioning of down-stream neural circuits (Dhruv and Carandini, 2014; Griffanti et al., 2014; Haak and Mesik, 2016; Haak et al., 2014, 2015; Schwartz et al., 2007; Seriès et al., 2009; Wandell and Smirnakis, 2009; Webster, 2015), which would require higher levels to update their interpretation of the neural code. Thus, it is important to maintain an appropriate balance between stability and plasticity.

Whether and how the balance between stability and plasticity varies across the brain is an important open question. It is thought that plasticity increases up cortical processing hierarchies (Fine, 2007; Haak et al., 2015; Huxlin, 2008), which would be consistent with the hypothesis that lower level plastic changes are more costly because more dependent processing stages would have to update their interpretation of the neural code (Haak et al., 2015). In addition, higher order areas appear more experience dependent and they contain neurons that prefer stimuli that have been frequently encountered or that are behaviourally relevant (Fine, 2007; Rolls and Tovee, 1995; Yang and Maunsell, 2004; Young and Yamane, 1992). The idea that plasticity is increased at higher levels of processing also agrees with observations that learning scales with task complexity, with lower-level tasks supported by lower-order areas showing less learning, and that neural changes related to learning appear larger in higher-order areas (Fine, 2007; Fine and Jacobs, 2002). However, so far, no direct quantitative comparisons between lower-and higher-level plasticity have been made. Here, we addressed this issue for the human cortical visual system.

Determining whether and how plasticity varies across the visual processing hierarchy requires a quantifiable measurement process. One possible approach would be to characterise the configuration of the neural circuitry at one point in time and then determine how much it changed at a later point. This approach requires a longitudinal study design. Another, equally valid approach is to assess the current state of configuration with respect to a state where the configuration was free of environmental influence. Such a state of zero-change corresponds to the configuration of neural circuits that is completely determined by the genetic blueprint, and hence plasticity can be quantified as the complement of the amount of phenotypic variance that can be explained by genetic factors. The amount of phenotypic variance that can be explained by genetic factors is known as heritability, which can be estimated under a twin study design. In the present work, we adopted the latter approach, as it allowed us to gauge the totality of plastic changes that occurred across the entire lifespan up to the time of measurement, and because it enabled answering our research question based on the publicly available neuroimaging data of the WU-Minn Human Connectome Project (Van Essen et al., 2013).

There are several possible phenotypes that can be assessed. For instance, plastic changes might be assessed in terms of stimulus and/or task-related neural responses. However, it is often unclear what stimulus or task should be used to target specific processing levels, and under external stimulation and/or a task it would be difficult to distinguish true variations in plasticity from constant plasticity with differences in expression due to limitations imposed by neuronal response properties specific to each processing level (Huxlin, 2008). In addition, although for primary processing nodes (e.g. the retina of the eye) plasticity may be defined as a change in response to external stimulation, for higher-order processing nodes it ought to be defined as a change in response to a change in the signals these nodes receive from lower-level processing stages (i.e., because response changes at higher-levels could be a manifestation of lower-level plastic changes). Plastic changes might also be assessed in terms of anatomical features. However, plasticity is not limited to anatomical changes, as evidenced by, for instance, adaptive neural tuning changes in response temporarily altered image statistics (Webster, 2015) and the existence of long-term potentiation (LTP) to facilitate learning and memory by synaptic strengthening (Cooke and Bear, 2010). We therefore chose functional connectivity as our phenotype of interest, which can be determined using human neuroimaging for every visual area in the absence of a task and/or external stimulation, and—in contrast to anatomical connectivity—is sensitive to both structural and functional plastic changes.

## Results

To quantify the balance between plasticity and stability across human visual cortex, we determined the functional connectivity between 48 cortical visual areas based on the publicly available resting-state functional MRI (rfMRI) data of twins provided by the WU-Minn Human Connectome Project (HCP) (Van Essen et al., 2013; Smith et al., 2013). Our sample included only those subjects whose twin-status was genetically confirmed, and who had completed all rfMRI sessions. Thus, our sample consisted of 123 monozygotic (MZ) and 67 dizygotic (DZ) twin-pairs (380 subjects in total).

Visual areas were delineated using a publicly available atlas of human retinotopic cortex (Wang et al., 2014), and we extracted the average rfMRI time-series from each of them. For each subject, we regressed out the effects of head-movement and computed functional connectivity between each area-pair as the correlation between the residual time-series. The statistical significance of each connection was assessed by block-permutation (Winkler et al., 2015), and all estimates whose FDR corrected *p*-value exceeded .05 were excluded from further analysis. The heritability (*h*^2^) of the functional connectivity between all possible area-pairs was estimated with covariates age, sex, MR reconstruction software version, and a summary measure of each subject’s head motion during scanning (i.e., mean frame-wise displacement). Figure 1A shows the estimates for each statistically significant (*p* < .05, FDR corrected) connection.

**Figure 1.**
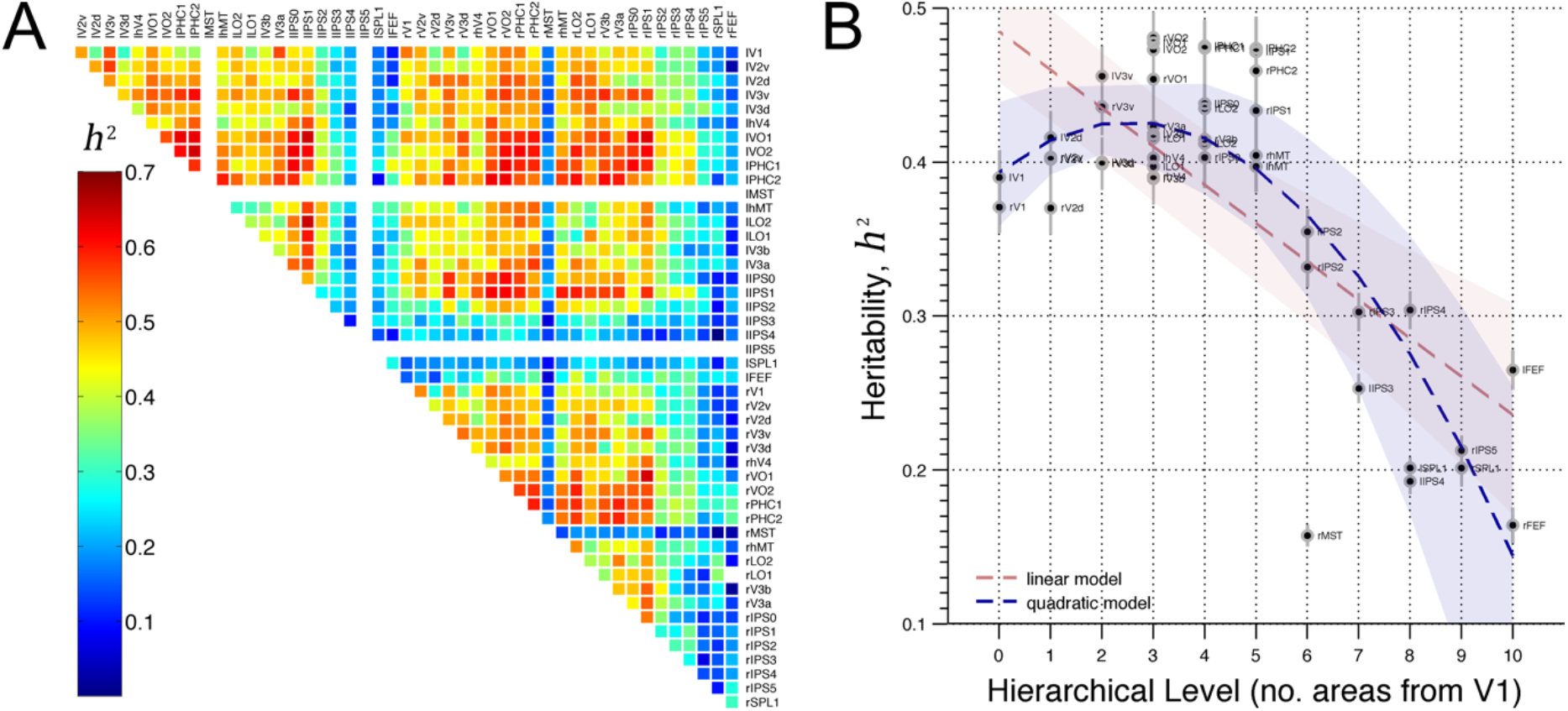
Heritability of functional connectivity across human visual cortex. **(A)** Heritability estimates for all significant (*p* < .05, FWE corrected) functional connections. Visual area names are abbreviated according to Wang et al. (2015). **(B)** Average heritability of functional connectivity as a function of hierarchical level. Grey error-bars indicate the SEM. Dashed red and blue lines and similarly coloured shaded areas indicate linear and quadratic model fits and the bootstrapped 95% confidence intervals, respectively.

We first ascertained that the heritability estimates could not be explained by possible confounding effects. For instance, visual areas differ in size and the temporal signal-to-noise ratio also varied across the occipital lobe. This could have led to differences in measurement error, which could in turn have influenced the heritability estimates. However, this was not the case, because the test-retest reliability of the connectivity estimates did not covary with heritability (Mantel test: *R*^2^ = .01, *p* = .33). In addition, atlas-based area definitions are not expected to fit equally perfectly across subjects, which could have influenced the heritability estimates because this misfit is likely heritable in its own right (Ge et al., 2016). However, this was also not the case, because the heritability estimates were unrelated to the expected precision of area definition (*R*^2^ = .02, *F*_1,44_ = 1.07, *p* = .31)—see Methods for details.

Next, we asked whether the heritability of each area’s connectivity ‘fingerprint’ was related its hierarchical level. To this end, we computed for each visual area the average heritability across all of its connections. As in previous work (Haak and Beckmann, 2018), we determined each area’s hierarchical level as the number of visual areas between that area and area V1. Ideally, the hierarchical level of a visual area is determined by assessing the laminar origin and termination patterns of their connections (Felleman and Van Essen, 1991). However, such information is not yet available for humans and the present approximation corresponds well to a data-driven estimation of hierarchical level based on multidimensional scaling (Haak and Beckmann, 2018), which also correctly determined hierarchical level from a matrix based on the laminar origin and termination patterns in the macaque (Young, 1992).

Figure 1B shows the heritability of each area’s connectivity fingerprint as a function of hierarchical level. To test if heritability was significantly related to hierarchical level, we performed linear and quadratic regression analyses. Whilst both models indicated a highly significant relation between heritability and hierarchical level (linear: *R*^2^ = .49, *F*_1,44_ = 43, *p* = 5.1 ×10^-8^; quadratic: *R*^2^ = .67, *F*_2,43_ = 44.1, *p* = 3.8×10^-11^), the quadratic model fitted the data better than the linear model (*F*_1,43_ = 23.3, *p* = 1.8×10^-5^) because heritability increased significantly from V1 to V3 (*t*_8_ = 2.38, *p* = .04). Thus, plasticity (i.e. the complement of heritability) is indeed greater at higher versus lower levels of visual processing, but it does not increase monotonically up the visual processing hierarchy.

## Discussion

We investigated whether and how the plasticity of functional connectivity varies across the cortical visual hierarchy. To this end, we leveraged the fact that plasticity can be quantified as the complement of heritability. That is, we quantified plasticity as the amount of deviation from the genetic blueprint. The results indicate a non-monotonic relationship between plasticity and hierarchical level, such that plasticity decreases from V1 to V3 and then increases further up the visual hierarchy.

The observation that plasticity decreases from V1 to V3 indicates that the first stages of cortical visual processing are not the most stable. This challenges theory that the amount of plasticity depends on a neural circuit’s hierarchical level because plastic changes at one level cause cascading effects downstream where higher level circuits need to update their interpretation of the new neural code (Haak and Mesik, 2016; Haak et al., 2014, 2015; Schwartz et al., 2007; Wandell and Smirnakis, 2009). This “coding catastrophe” confers lower costs to plasticity at higher levels of the processing hierarchy, because higher levels have fewer downstream dependents. If these costs critically determined the amount of plasticity along cortical processing hierarchies, plasticity should monotonically increase up the visual hierarchy, and the first stage of cortical processing (V1) should be the most stable. However, the present data suggest that this is not the case.

Why plasticity is distributed this way is an important open question. One possibility is that plasticity decreases along the ventral occipital surface, whereas it increases up the lateral and dorsal occipital surfaces. Indeed, areas along the ventral occipital surface exhibited greater heritability than areas on the lateral and dorsal occipital surfaces (Figure 1B), and previous work also noted differences in ventral versus dorsal visual plasticity (Braddick and Atkinson, 2011; Huxlin, 2008). However, while it might be reasonable to speculate that higher level dorsal visual areas are more plastic because they are increasingly involved in interacting with the environment, it is more difficult to explain why plasticity would decrease along the ventral occipital surface.

Another possibility is that the increasing heritability from V1 to V3 reflects decreasing transient adaptive changes in response to recent visual experience, whereas the overall decline in heritability reflects a rise in more permanent plastic changes due to learning. Indeed, different types of plasticity may be distinctly expressed at different cortical locations, and it is therefore possible that short-term plasticity is most pronounced in V1 and then decreases up the visual hierarchy, while long-term plasticity is increasingly pronounced in higher level visual areas (see Figure 2). This account agrees with both a lack of long-term V1 plasticity after retinal damage (Baseler et al., 2011; Sereno, 2005; Smirnakis et al., 2005; Wandell and Smirnakis, 2009) and the large body of evidence of short-term adaptation (Webster, 2015) in early visual cortex.

**Figure 2.**
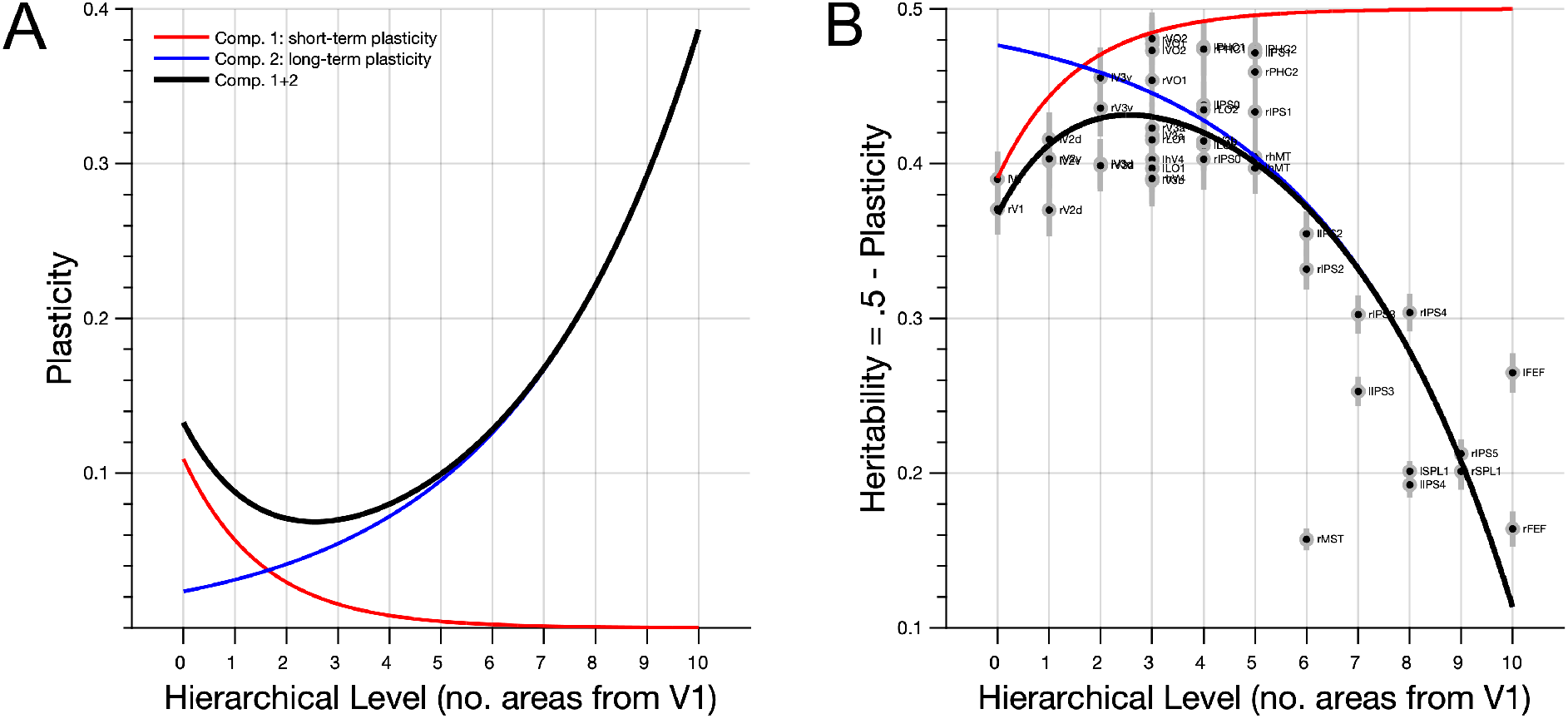
Theoretical model of the non-monotonic relationship between heritability and hierarchical level based on distinct distributions of short-term and long-term plasticity. **(A)** The non-monotonic relationship between heritability and hierarchical level can be modelled as the complement of the sum of transient short-term plasticity and sustained long-term plasticity components. Under this model, short-term plasticity (red) is relatively weak and decreases exponentially from low to high levels of visual processing, whereas long-term plasticity (blue) increases exponentially to become relatively strong in high-level visual cortex. **(B)** Under the assumption that the average heritability estimates shown in Figure 1B peak at .5 (due to e.g. measurement noise common to all hierarchical levels of processing), the complement of the sum of both components (black) predicts the observed heritability estimates very well (*R*^2^ = .73).

Importantly, these accounts are not mutually exclusive because visual processing along the ventral occipital surface might be primarily governed by short-term plastic changes— indeed, the red line in Figure 2B adheres closely to the heritability estimates for the ventral visual areas—whereas visual processing along the lateral and dorsal occipital surfaces might be governed by the sum of short and long-term plasticity (thick black line in Figure 2B).

By quantifying plasticity as the complement of heritability, our measure covers the totality of all possible changes with respect to the genetic blueprint across the entire lifespan up to the time of measurement. This means that we cannot comment on whether these changes occurred during childhood or adulthood. It is further possible that some of the changes we attribute to plasticity are in fact non-plastic changes that occurred due to ageing, injury or disease. However, because our sample included only healthy young adults in whom the visual system should be fully developed but not yet aged (Haak, 2018), the contribution of such non-plastic changes to the overall amount of change should be negligible.

In conclusion, our results indicate that the notion of increased plasticity at higher levels of cortical processing is an over-simplification. Rather, they suggest that there are different types of plasticity (e.g., short- and long-term plasticity), the expression of which varies distinctly across the cortical visual system. This principle may apply across the sensory modalities and species. While our results pertain principally to plastic changes under normal conditions, they may also have clinical relevance when used to gauge how much plasticity can be expected in response to focal brain injuries and neurological disease.

## Methods

### Dataset and pre-processing

The dataset comprised subjects of the S1200 WU-Minn HCP (Van Essen et al., 2013) data-release who completed all of the four rfMRI runs and whose twin-status was confirmed by genetic testing. The dataset included 123 monozygotic (MZ) and 67 dizygotic (DZ) twin-pairs. Their 2^3^mm^3^, multiband accelerated (x8) 3T rfMRI data with a TR of .72s were pre-processed as detailed in (Smith et al., 2013), which involved rigorous data cleaning using the FIX artefact removal procedure (Griffanti et al., 2014; Salimi-Khorshidi et al., 2014). For the present work, we additionally smoothed the images using a 3mm FWHM Gaussian kernel.

### Regions-of-interest definition

Visual areas were defined using a probabilistic atlas (Wang et al., 2014). The atlas provides both full probability maps and maximum probability maps (i.e., the most probable area label for any given point) in MNI space. We used the latter for region-of-interest definition and down-sampled the region definitions from 1 mm to 2 mm isotropic resolution using nearest-neighbour interpolation. Furthermore, a single V1 region was defined by combining the maximum probability maps labelled V1d and V1v in each hemisphere. As such, the number of regions-of-interest that were used in subsequent analysis steps was 48 (24 in each cerebral hemisphere).

### Functional connectivity analysis

We determined the functional connectivity between all possible area-pairs at the individual level by (1) extracting the average time-series from each region-of-interest, (2) regressing out the motion realignment parameters and their first derivatives, and (3) computing the (Fisher’s r-to-z transformed) correlation between the residuals. Next, we determined the group-level statistical significance of the functional connectivity estimates using a block-permutation test (Winkler et al., 2015) to account for the family-structure in the dataset. The ensuing p-values were corrected for multiple comparisons using a false-discovery rate (FDR) approach, and all connections whose FDR-corrected p-value exceeded .05 were excluded from further analysis.

### Heritability analysis

The heritability of each connection was estimated using the freely available SOLAR software package (Almasy and Blangero, 1998). Heritability, which is defined as the portion of phenotypic variance accounted for by the total additive genetic variance (i.e., 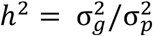), was assessed with simultaneous estimation for the effects of covariates age, sex, a measure of each subject’s head motion during scanning (mean frame-wise displacement), and MRI reconstruction software version^1^. SOLAR estimates the variances 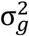 and 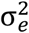 by comparing the observed phenotypic covariance with the covariance predicted by kinship (i.e., 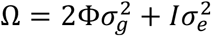), where Φ is the kinship matrix), and determines the statistical significance of the heritability estimates by comparing the log-likelihood of the model in which 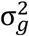 is constrained to be zero (*L*_0_) to the log-likelihood of the model in which 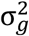 is estimated (*L*_e_). This is done using the test-statistic 2(*L*_e_ -*L*_0_) which is asymptotically distributed as a 50:50 mixture of a 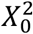 (point mass) and a 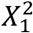 distribution. Prior to analysis, the functional connectivity estimates were subjected to the inverse normal transformation to ensure that the residual kurtosis (i.e. the kurtosis after the covariate influences have been removed) was within normal range.

Given that both the size and temporal signal-to-noise ratio (tSNR) of the visual areas varied across the occipital lobe, it was important to rule out that the heritability estimates were biased by measurement error. Therefore, we leveraged the fact that each subject was scanned on two different session days and re-computed the functional connectivity matrix for each session day to determine the test-retest reliability. If the heritability estimates co-varied with measurement error, this would result in a correlation between the test-retest reliability across connections and heritability. Thus, we computed the intra-class correlation coefficient (ICC_3,1_) (Shrout and Fleiss, 1979; Zuo et al., 2010) for each connection and then determined the variance in the heritability matrix (Fig 1A) that could be explained by the ICC matrix. We used a non-parametric permutation approach to test for statistical significance (i.e., a Mantel test with 5000 permutations), because the values within either matrix are not independent.

In addition, we ascertained that the heritability estimates shown in Fig 1B were not related to the expected precision of area definition. This was important because atlas-based area definitions are not expected to fit equally perfectly across subjects, which could have influenced the heritability estimates. To verify that this was not the case, we regressed the likelihood that cortical points in a given area would be classified as part of that area in unseen subjects (i.e., the mean of the corresponding probability map masked with the atlas definition of that area) onto the heritability estimates (averaged across all of an area’s connections because we only have one estimate of area definition precision per area), and determined statistical significance using an F-test.

### Regression analyses

To determine the relation between the heritability of functional connectivity and hierarchical level, we first summarized the heritability of each area’s connections by averaging the heritability estimates across all of that area’s connections. By doing so, we effectively assess the heritability of an area’s entire functional connectivity profile with respect to the rest of visual cortex (i.e., its functional connectivity fingerprint), with the added benefit that there is no need to further correct for effects of distance because the average distance from one area to all other areas is equal for all areas. As in previous work (Haak and Beckmann, 2018), we determined the hierarchical level of each visual area by constructing a nearest-neighbour graph with edges between areas only if they are direct neighbours. We then used Dijkstra’s algorithm to determine the shortest path through this graph from each visual area to V1. The length of this shortest path (i.e., the number of areas that need to be visited before V1 can be reached) was our measure of hierarchical level.

We considered two possible relationships between heritability and hierarchical level: (1) a linear model (i.e., 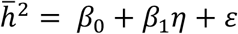, where *η* represents hierarchical level) in accordance with the hypothesis that heritability decreases monotonically up the visual hierarchy, and (2) a quadratic model (i.e., 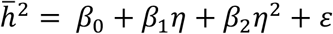), because visual inspection of the data strongly suggested that the heritability of early visual connectivity was much lower than might be expected had the relation been a strictly monotonic linear decrease. The significance of the two models was assessed by F-tests and the two (nested) models were compared using a *F*-ratio test. Finally, we determined the relationship between heritability and hierarchical level within early visual cortex (V1, V2d, V2v, V3d, V3d in both hemispheres; 10 areas total) by linear regression, and assessed the statistical significance of the slope (*t*-test).

### Two-component model of plasticity

To model the observed relationship between heritability and hierarchical level, we assumed that heritability peaks at .5 (to account for unbiased measurement noise common to all stages of visual processing) and that total plasticity can be decomposed as the sum of two components: one of which is relatively weak and decreases exponentially with hierarchical level (i.e., *c*_1_ = *a*_1_· *e*^-*b*_1_*x*^, where *x* is hierarchical level), and one that is relatively strong and increases exponentially with hierarchical level (i.e., *c*_2_ = *a*_2_· *e*^*b*_2_*x*^). As such, the observed heritability is predicted by *h*^2^ = .5 – (*c*_2_ + *c*_2_). To Parameters *a*_1_, *b*_1_, *a*_2_ and *b*_2_ were estimated using robust non-linear least-squares regression.

## Footnotes

1 https://wiki.humanconnectome.org/display/PublicData/Ramifications+of+Image+Reconstruction+Version+Differences

## Acknowledgments

This work was supported by the Netherlands Organization for Scientific Research Veni Grant No. 016.Veni.171.068 (to KVH), and Vidi Grant No. 864-12-003 (to CFB). Data were provided by the Human Connectome Project, WU-Minn Consortium (Principal Investigators: David Van Essen and Kamil Ugurbil; 1U54MH091657) funded by the 16 NIH Institutes and Centers that support the NIH Blueprint for Neuroscience Research; and by the McDonnell Center for Systems Neuroscience at Washington University.

